# Heterogeneous Network Edge Prediction: A Data Integration Approach to Prioritize Disease-Associated Genes

**DOI:** 10.1101/011569

**Authors:** Daniel S. Himmelstein, Sergio E. Baranzini

## Abstract

The first decade of Genome Wide Association Studies (GWAS) has uncovered a wealth of disease-associated variants. Two important derivations will be the translation of this information into a multiscale understanding of pathogenic variants, and leveraging existing data to increase the power of existing and future studies through prioritization. We explore edge prediction on heterogeneous networks—graphs with multiple node and edge types—for accomplishing both tasks. First we constructed a network with 18 node types—genes, diseases, tissues, pathophysiologies, and 14 MSigDB (molecular signatures database) collections—and 19 edge types from high-throughput publicly-available resources. From this network composed of 40,343 nodes and 1,608,168 edges, we extracted features that describe the topology between specific genes and diseases. Next, we trained a model from GWAS associations and predicted the probability of association between each protein-coding gene and each of 29 well-studied complex diseases. The model, which achieved 132-fold enrichment in precision at 10% recall, outperformed any individual domain, highlighting the benefit of integrative approaches. We identified pleiotropy, transcriptional signatures of perturbations, pathways, and protein interactions as fundamental mechanisms explaining pathogenesis. Our method successfully predicted the results (with AUROC = 0.79) from a withheld multiple sclerosis (MS) GWAS despite starting with only 13 previously associated genes. Finally, we combined our network predictions with statistical evidence of association to propose four novel MS genes, three of which (*JAK2*, *REL*, *RUNX3*) validated on the masked GWAS. Furthermore, our predictions provide biological support highlighting *REL* as the causal gene within its gene-rich locus. Users can browse all predictions online (http://het.io). Heterogeneous network edge prediction effectively prioritized genetic associations and provides a powerful new approach for data integration across multiple domains.

**Author Summary:** For complex human diseases, identifying the genes harboring susceptibility variants has taken on medical importance. Disease-associated genes provide clues for elucidating disease etiology, predicting disease risk, and highlighting therapeutic targets. Here, we develop a method to predict whether a given gene and disease are associated. To capture the multitude of biological entities underlying pathogenesis, we constructed a heterogeneous network, containing multiple node and edge types. We built on a technique developed for social network analysis, which embraces disparate sources of data to make predictions from heterogeneous networks. Using the compendium of associations from genome-wide studies, we learned the influential mechanisms underlying pathogenesis. Our findings provide a novel perspective about the existence of pervasive pleiotropy across complex diseases. Furthermore, we suggest transcriptional signatures of perturbations are an underutilized resource amongst prioritization approaches. For multiple sclerosis, we demonstrated our ability to prioritize future studies and discover novel susceptibility genes. Researchers can use these predictions to increase the statistical power of their studies, to suggest the causal genes from a set of candidates, or to generate evidence-based experimental hypothesis.

## Introduction

In the last decade, genome-wide association studies (GWAS) have been established as the main strategy to map genetic susceptibility in dozens of complex diseases and phenotypes. Despite the undeniable success of this approach, researchers are now confronted with the challenge of maximizing the scientific contribution of existing GWAS datasets, whose undertakings represented a substantial investment of human and monetary resources from the community at large.

A central assumption in GWAS is that every region in the genome (and hence every gene) is a-priori equally likely to be associated with the phenotype in question. As a result, small effect sizes and multiple comparisons limit the pace of discovery. However, rational prioritization approaches may afford an increase in study power while avoiding the constraints and expense related to expanded sampling. One such a way forward is the current trend on analyzing the combined contribution of susceptibility variants in the context of biological pathways, rather than single SNPs [1, 2]. A less explored but potentially revealing strategy is the integration of diverse sources of data to build more accurate and comprehensive models of disease susceptibility.

Several strategies have been attempted to identify the mechanisms underlying pathogenesis and use these insights to prioritize genes for genetic association analyses. Gene-set enrichment analyses identify prevalent biological functions amongst genes contained in disease-associated loci [3]. Gene network approaches search for neighborhoods of genes where disease-associated loci aggregate [4]. Literature mining techniques aim to chronicle the relatedness of genes to identify a subset of highly-related associated genes [5]. These strategies generally rely on user-provided loci as the sole input and do not incorporate broader disease-specific knowledge. Typically, the proportion of genome-wide significant discoveries in a given GWAS is low, thus leaving little high-confidence signal for seed-based approaches to build from. To overcome this limitation, here we aimed at characterizing the ability of various information domains to identify pathogenic variants across the entire compendium of complex disease associations. Using this multiscale approach, we developed a framework to prioritize both existing and future GWAS analyses and highlight candidate genes for further analysis.

To approach this problem, we resorted to a method that integrated diverse information domains naturally. Heterogeneous networks are a class of networks which contain multiple types of entities (nodes) and relationships (edges), and provide a data structure capable of expressing diversity in an intuitive and scalable fashion. However, current techniques available for network analysis have been developed for homogeneous networks and are not directly extensible to heterogenous networks. Furthermore, research into heterogeneous network analysis is in its early stages [6]. One of the few existing methods for predicting edges on heterogeneous networks was developed by researchers studying social sciences to predict future coauthorship [7]. In this work, we extended this methodology to predict the probability that an association between a gene and disease exists.

## Results

### Constructing a heterogeneous network to integrate diverse information domains

Using publicly-available databases and standardized vocabularies, we constructed a heterogeneous network with 40,343 nodes and 1,608,168 edges (Figure 1). Databases were selected based on quality, reusability, and throughput. The network was designed to encode entities and relationships relevant to pathogenesis. The network contained 18 node types (metanodes) and 19 edge types (metaedges), displayed in Figure S2A. Entities represented by metanodes consisted of diseases, genes, tissues, patho-physiologies, and gene sets for 14 MSigDB collections including pathways, perturbation signatures, motifs, and Gene Ontology domains (Table 1). Relationships represented by metaedges consisted of gene-disease association, disease pathophysiology, disease localization, tissue-specific gene expression, protein interaction, and gene-set membership for each MSigDB collection (Table 2).

**Figure 1.**
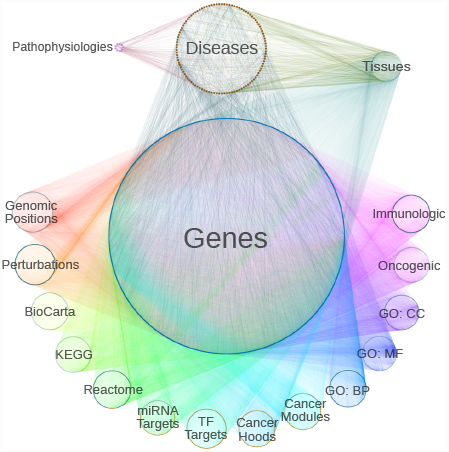
Heterogeneous network integrates diverse information domains. We constructed a heterogeneous network with 18 metanodes (denoted with labels) and 19 metaedges (denoted by color). For each metanode, nodes are laid out circularly. Incorporating type information adds structure to a network which would otherwise appear as an undecipherable agglomeration of 40,343 nodes and 1,608,168 edges.

**Table 1.**
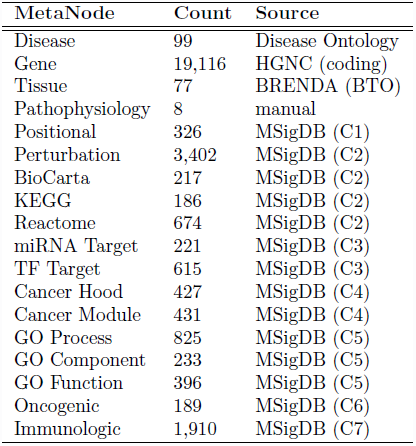
Metanodes. The kind, number of corresponding nodes, and data source for each type of node.

**Table 2.** Metaedges. The kind, number of corresponding edges, and data source for each type of edge.

| MetaEdge | Count | Source |
| --- | --- | --- |
| Disease - association - Gene | 938 | GWAS Catalog |
| Disease - membership - Pathophysiology | 90 | manual |
| Disease - localization - Tissue | 1,086 | CoPub 5.0 |
| Gene - expression - Tissue | 251,366 | GNF BodyMap |
| Gene - interaction - Gene | 97,938 | iRefIndex |
| Gene - membership - Positional | 18,343 | MSigDB (C1) |
| Gene - membership - Perturbation | 366,211 | MSigDB (C2) |
| Gene - membership - BioCarta | 4,456 | MSigDB (C2) |
| Gene - membership - KEGG | 12,656 | MSigDB (C2) |
| Gene - membership - Reactome | 35,597 | MSigDB (C2) |
| Gene - membership - miRNA Target | 33,455 | MSigDB (C3) |
| Gene - membership - TF Target | 161,258 | MSigDB (C3) |
| Gene - membership - Cancer Hood | 41,913 | MSigDB (C4) |
| Gene - membership - Cancer Module | 48,220 | MSigDB (C4) |
| Gene - membership - GO Process | 75,155 | MSigDB (C5) |
| Gene - membership - GO Component | 34,880 | MSigDB (C5) |
| Gene - membership - GO Function | 23,578 | MSigDB (C5) |
| Gene - membership - Oncogenic | 30,166 | MSigDB (C6) |
| Gene - membership - Immunologic | 370,862 | MSigDB (C7) |

Gene-disease associations were extracted from the GWAS Catalog [8] by overlapping associations into disease-specific loci. Loci were classified as low or high-confidence based on p-value and sample size of the corresponding GWAS. When possible, for each loci, the most-commonly reported gene across studies was designated as primary and subsequently considered responsible for the association. Additional genes reported for the loci were considered secondary. Only high-confidence primary associations were included in the network yielding 938 associations between 99 diseases and 711 genes (Figure S1 visualizes a subset of these associations).

### Features quantify the network topology between a gene and disease

To describe the network topology connecting a specific gene and disease, we computed 24 features, each describing a different aspect of connectivity. Each feature corresponds to a type of path (metapath) originating in a given source gene and terminating in a given target disease. The biological interpretation of a feature derives from its metapath (Table S1), and features simply quantify the prevalence of a specific metapath between any gene-disease pair. To quantify metapath prevalence, we adapted an existing method originally developed for social network analysis (*PathPredict*) [7], and developed a new metric called degree-weighted path count (*DWPC*, Figure S2D), which we employed in all but two features. The *DWPC* downweights paths through high-degree nodes when computing metapath prevalence. The strength of downweighting depends on a single parameter (*w*), which we optimized to *w* = 0.4 and that outperformed the top metric resulting from *PathPredict* (Figure S3A) [7]. Two non-*DWPC* features were included to assess the pleiotropy of the source gene and the polygenicity of the target disease. Referred to as ‘path count’ features, they respectively equal the number of diseases associated with the source gene and the number of genes associated with the target disease. For all features, paths with duplicate nodes were excluded, and, if present, the association edge between the source gene and target disease was masked.

### Machine learning approach to predict the probability of association of gene-disease pairs

Further analysis focused on the 29 diseases with at least ten associated genes (Table 3). The 698 high-confidence primary associations of these 29 diseases were considered positives—gene-disease pairs with positive experimental relationships (as defined in Methods, Figure S1). The remaining 551,823 (i.e. unassociated) gene-disease pairs were considered negatives. Low-confidence or secondary associations were excluded from either set. We partitioned gene-disease pairs into training (75%) and testing (25%) sets and created a training network with the testing associations removed.

**Table 3.** Diseases. Associations were predicted for 29 diseases with at least 10 positives. For these diseases, the number of high-confidence primary (HC-P), high-confidence secondary (HC-S), low-confidence primary (LC-P), and low-confidence secondary associations (LC-S) that were extracted from the GWAS Catalog is indicated.

| Disease | Pathophysiology | HC-P | HC-S | LC-P | LC-S |
| --- | --- | --- | --- | --- | --- |
| Crohn's disease | immunologic | 67 | 179 | 4 | 2 |
| multiple sclerosis | immunologic | 50 | 43 | 38 | 29 |
| type 2 diabetes mellitus | immunologic | 49 | 49 | 20 | 15 |
| breast carcinoma | neoplastic | 43 | 65 | 2 | 6 |
| ulcerative colitis | immunologic | 40 | 96 | 2 | 3 |
| prostate carcinoma | neoplastic | 34 | 202 | 3 | 4 |
| type 1 diabetes mellitus | immunologic | 33 | 56 | 9 | 6 |
| rheumatoid arthritis | immunologic | 30 | 27 | 20 | 11 |
| coronary artery disease | metabolic | 29 | 43 | 15 | 9 |
| obesity | metabolic | 28 | 22 | 34 | 18 |
| celiac disease | immunologic | 24 | 32 | 9 | 8 |
| systemic lupus erythematosus | immunologic | 22 | 35 | 14 | 8 |
| refractive error | degenerative | 21 | 11 | 2 | 1 |
| primary biliary cirrhosis | immunologic | 20 | 16 | 2 | 0 |
| vitiligo | immunologic | 20 | 27 | 4 | 0 |
| age related macular degeneration | degenerative | 18 | 30 | 11 | 18 |
| metabolic syndrome X | metabolic | 17 | 11 | 1 | 0 |
| asthma | immunologic | 17 | 23 | 13 | 4 |
| psoriasis | immunologic | 16 | 14 | 5 | 5 |
| schizophrenia | psychiatric | 15 | 27 | 20 | 13 |
| chronic lymphocytic leukemia | neoplastic | 14 | 16 | 3 | 4 |
| migraine | unspecific | 13 | 15 | 38 | 58 |
| Alzheimer's disease | degenerative | 12 | 11 | 27 | 18 |
| Graves' disease | immunologic | 12 | 15 | 1 | 1 |
| Parkinson's disease | degenerative | 12 | 21 | 8 | 13 |
| atopic dermatitis | immunologic | 11 | 15 | 5 | 1 |
| bipolar disorder | psychiatric | 11 | 34 | 26 | 74 |
| lung carcinoma | neoplastic | 10 | 14 | 6 | 6 |
| ankylosing spondylitis | immunologic | 10 | 5 | 6 | 6 |

To learn the importance of each feature and model the probability of association of a given gene-disease pair, we used regularized logistic regression which is designed to prevent overfitting and accurately estimate regression coefficients when models include many features. Elastic net regression is a regression method that balances two regularization techniques: ridge (which performs coefficient shrinkage) and lasso (which performs coefficient shrinkage and variable selection) [9]. On the training set, we optimized the elastic net mixing parameter, a single parameter behind the *DWPC* metric, and two edge-inclusion thresholds (Figure S3). While cross-validated performance was similar across elastic net mixing parameters, ridge demonstrated the greatest consistency (Figure S3A), and thus we proceeded with logistic ridge regression as the primary model for predictions.

### Method prioritizes associations withheld for testing

We extracted network-based features for gene-disease pairs from the training network and modeled the training set. We next evaluated performance on the 25% of gene-disease pairs (175 positives, 137,956 negatives) withheld for testing. Our predictions achieved an area under the ROC curve (AUROC) of 0.83 (Figure 2A) demonstrating an excellent performance in retrieving hidden associations. Importantly, we did not observe any significant degradation of performance from training to testing (Figure 2A), indicating that our disciplined regularization approach avoided overfitting and that predictions for associations included in the network were not biased by their presence in the network. Furthermore, we observed that at 10% recall (the classification threshold where 10% of true positives were predicted as positives), our predictions achieved 16.7% precision (the proportion of predicted positives that were correct). Since the prevalence of positives in our dataset was 0.13%, the observed precision represents a 132-fold enrichment over the expected probability under a uniform distribution of priors (as in GWAS).

**Figure 2.**
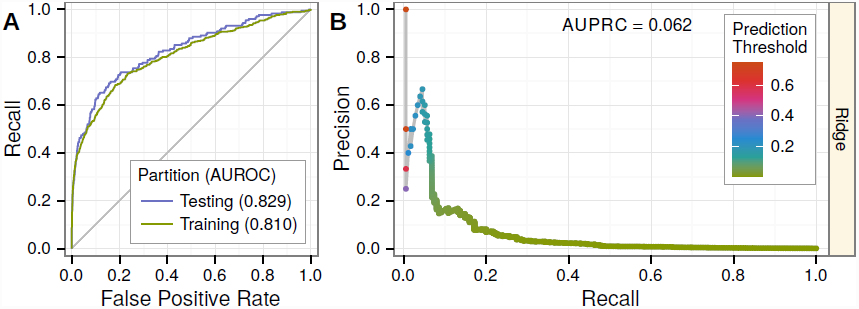
Predicting associations withheld for testing. Performance was evaluated on 25% of gene-disease pairs withheld for testing. A) Testing and training ROC curves. At top prediction thresholds, associated gene-disease pairs are recalled at a much higher rate than unassociated pairs are incorrectly classified as positives. The testing area under the curve (AUROC) is slightly greater than the training AUROC, demonstrating the method’s lack of overfitting. Performance greatly exceeds random denoted by gray line. B) The precision-recall curve showing performance in the context of the low prevalence of associated gene-disease pairs (0.13%). Nevertheless, at top prediction thresholds, a high percentage of pairs classified as positives are truly associated. Prediction thresholds, shown as points and colored by value, align with the observed precision at that threshold.

### Predicting associations on the complete network

As a next step in our analysis, we recomputed features on the complete network, which now included the previously withheld testing associations. On all positives and negatives, we fit a ridge model (the primary model for predictions) and a lasso model (for comparison). Standardized coefficients (Figure 3) indicate the effect attributed to each feature by the models. The lasso highlighted features that captured pleiotropy (4 features), pathways (2), transcriptional signatures of perturbations (1) and protein interactions (1). Despite the parsimony of the lasso, performance was similar between models with training AUROCs of 0.83 (ridge) and 0.82 (lasso). However, since multiple features from a correlated group may be causal, the lasso model risks oversimplifying. Ridge regression disperses an effect across a correlated group of features, providing users greater flexibility when interpreting predictions. From the ridge model, we predicted the probability that each protein-coding gene was associated with each analyzed disease and built a webapp to display the predictions (http://het.io/disease-genes/browse).

**Figure 3.**
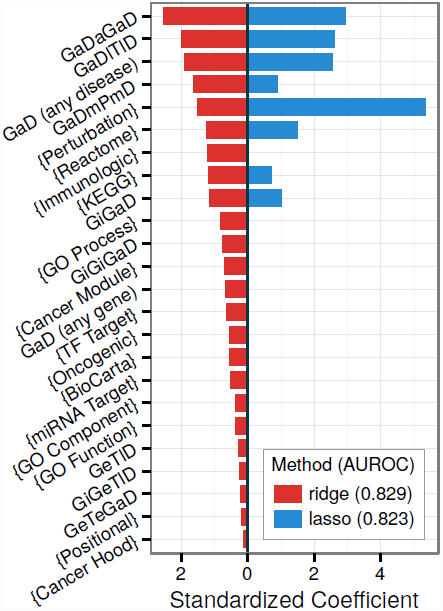
Feature selection identifies a parsimonious yet predictive model. Ridge and lasso models were fit from the complete network. The resulting standardized coefficients (x-axis) are plotted for each feature (y-axis). Brackets indicate features from MSigDB-traversing metapaths (*Gm*{}*mGaD*). The ridge model disperses effects amongst features whereas the lasso concentrates effects. The lasso identifies an 8-feature model with minimal performance loss compared to the ridge model. Besides *KEGG*, gene-set based features were largely captured by *Perturbations*. The lasso retains several measures of pleiotropy as well as the one-step interactome feature (*GiGaD*).

### Degree-preserving network permutations highlight the importance of edge-specificity for top predictions and ten features

Using Markov chain randomized edge-swaps, we created 5 permuted networks. Since metaedge-specific node degree is preserved, features extracted from the permuted network retain unspecific effects. These effects include general measures a disease’s polygenicity and a gene’s pleiotropy, multifunctionality, and tissue-specificity. On the first permuted network, we partitioned associations into training and testing sets. Testing associations were masked from the network, features were computed, and a ridge model was fit on the training gene-disease pairs.

Compared to the unpermuted-network model, testing performance was noticeably inferior: the AUROC declined from 0.83 (Figure 2A) to 0.79 (Figure S4A) and the AUPRC (area under the precision-recall curve) declined from 0.06 (Figure 2B) to 0.02 (Figure S4B). We interpret the modest decline in AUROC but marked reduction in AUPRC as a direct consequence of the permutation’s particularly detrimental effect on top predictions (Figure S4C–D). In other words, edge-specificity was crucial for top predictions, while general effects gleaned from node degree performed reasonably well when ranking the entire spectrum of protein-coding genes for association. A commonly-overlooked finding is that the discriminatory ability of gene networks largely relies on node-degree rather than the edge-specificity [10]. However, we found that for top predictions—which are the only predictions considered by many applications—edge-specificity was critical.

Interestingly, predictions from the permuted-network model displayed a reduced dynamic range with none exceeding 4%, while predictions from the unpermuted-network model exceeded 75% (Figure S4D). Therefore, even though they achieve reasonable AUROC, the permuted-network predictions would have little utility as prior probabilities in a bayesian analysis where dynamic range is crucial. Furthermore, the signal present in permuted-network features was greatly diminished: few features survived the lasso’s selection resulting in an average lasso AUROC of 0.70 versus 0.80 for ridge (Figure S5). Permuting the network significantly reduced the predictiveness of features based on pleiotropy (2 features), protein interactions (2), transcriptional signatures of perturbations (1), tissue-specificity (1), pathways (3), and immunologic signatures (1) (Table S2). Six of the eight features selected by the lasso and eight of the top ten ridge features (ranked by standardized coefficients) were negatively affected by the permutation. Since our modeling technique preferentially selected/weighted features affected by permutation, we can infer that network components where edge-specificity matters underlie a large portion of predictions.

### Feature importance identifies the mechanisms underlying associations

We assessed the informativeness of each feature by calculating feature-specific AUROCs. Feature-specific AUROCs universally exceeded 0.5, indicating that network connectivity, regardless of type, positively discriminates associations. However, performance varied widely by feature and within feature from disease to disease (Figure 4). Top performing domains consisted of transcriptional signatures of perturbations (AUROC = 0.74), immunologic signatures (0.70), and pleiotropy (0.68, 0.67, 0.64, 0.63). Notably, the models greatly outperformed any individual feature, highlighting the importance of an integrative approach.

**Figure 4.**
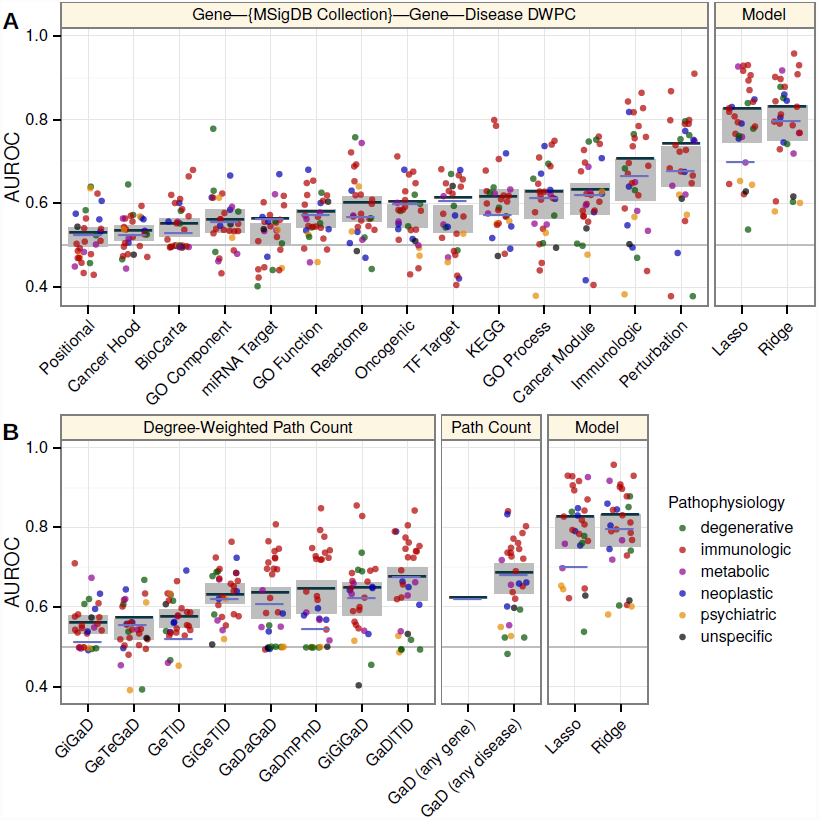
Decomposing performance shows the superiority of the integrative model and compares individual features. Disease, feature, and model-specific performance on the complete network. The AUROC (y-axis) was calculated for each classifier (x-axis). In addition to the ridge and lasso models (rightmost panels), each feature was considered as a classifier. Line segments show the classifier’s global performance (average performance across permuted networks shown in violet as opposed to dark grey). Points indicate disease-specific performance and are colored by the disease’s pathophysiology. Grey rectangles show the 95% confidence interval for mean disease-specific performance. A) Features from metapaths that traverse an MSigDB collection. B) Features from non-MSigDB-traversing metapaths. Metapaths are abbreviated using first letters of metanodes (uppercase, Table 1) and metaedges (lowercase, Table 2). Feature descriptions are provided in Table S1.

Features whose metapaths originate with an association (*GaD*) metaedge measure pleiotropy (Table S1). The four pleiotropic features were among the top performing features that did not rely on set-based gene categorization (Figure 4). Of the four features, *GaD (any disease)* had the highest AUROC despite its lack of disease-specificity, reflecting both the sparsity of disease-specific features and the existence of genetic overlap between seemingly disparate diseases. *GaDmPmD* and *GaDaGaD* performed best for immunologic diseases and were affected by permutation, indicating that genetic overlap was greatest between immunologic diseases. On the other hand, the performance of *GaDlTlD* did not decrease after permutation indicating disease colocalization was not a primary driver of genetic overlap.

We also observed that the lasso regression model discarded the majority of features with a minimal performance deficit, suggesting redundancy among features. Indeed, pairwise feature correlations showed moderate collinearity among features (Figure S6). Collinearity was especially pervasive with respect to the *Perturbations* feature, explaining its threefold increase in standardized coefficient in the lasso versus ridge model. The disappearance of all but one other MSigDB-based feature in the lasso model indicated that *Perturbations*—the feature traversing chemical and genetic transcriptional signatures of perturbations—exhausted meaningful gene-set characterization. In other words, the faulty molecular processes behind pathogenesis align with and are encapsulated by the processes perturbed by chemical and genetic modifications. The *Immunologic signatures* feature—traversing gene-sets characterizing “cell types, states, and perturbations within the immune system”—was highly predictive and correlated with *Perturbations*. As expected this feature performed best for diseases with an immune pathophysiology. The one well-performing neoplastic disease (Figure 4) was chronic lymphocytic leukemia, a hematologic cancer with a strong immune component [11]. Additionally, the performance of both the *Perturbation* and *Immunologic* features was affected by permutation indicating information beyond the extent of a gene’s multifunctionality was encoded.

Existing network-based gene-prioritization methods, frequently rely solely on protein-protein interactions. Our results supported incorporating protein interactions as the two interactome-based features were discriminatory (AUROCs = 0.65, 0.56) and affected by permutation. However, when compared to the integrative models or other top-performing features, performance of features that relied solely on the interactome was severely limited. Pathways, another founding resource for many approaches, proved important with *KEGG* selected by the lasso and all three pathway resources (AUROCs = 0.61 for *KEGG*, 0.60 for *Reactome*, 0.55 for *BioCarta*) affected by permutation. The *GeTlD* feature—measuring to what extent a gene is expressed in tissues affected by the disease in question—peaked in performance around AUROC = 0.58 (Figure S3B), was affected by permutation, and required no preexisting knowledge of associated genes. In other words, while approaches based on tissue-specificity may have limited predictive ability on their own, they are broadly applicable (i.e. less susceptible to knowledge bias) and provide orthogonal information that could enhance the overall performance of a model.

### Case study: prioritizing multiple sclerosis associations

The WTCCC2 multiple sclerosis (MS) GWAS tested 465,434 SNPs for 9,772 cases and 17,376 controls and identified over 50 independently associated loci [12]. Since the GWAS Catalog excludes targeted arrays (such as ImmunoChip), this study remains the largest MS GWAS in the Catalog. To evaluate our method’s ability to prioritize associations identified in a future study, we masked the WTCCC2 MS study from the GWAS Catalog and created a pre-WTCCC2 network. The number of high-confidence primary MS associations was thus reduced from 50 to 13, with the 37 novel genes identified by WTCCC2 available to evaluate performance. On the pre-WTCCC2 network, we extracted features, fit a ridge model, and predicted each gene’s probability of association with MS. Amongst all 18,993 potentially novel genes, the 37 WTCCC2 genes were ranked highly (AUROC = 0.79, Figure 5).

**Figure 5.**
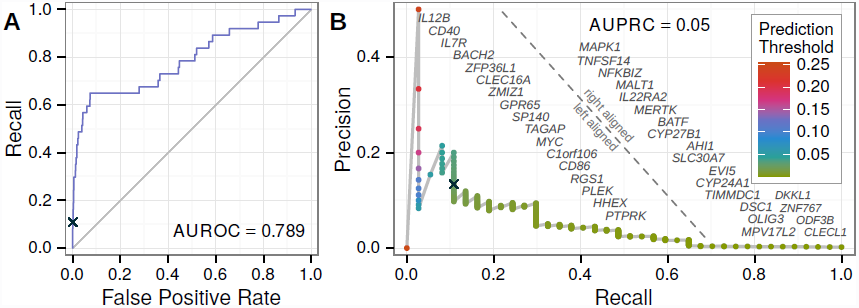
Prioritizing multiple sclerosis associations identified by a masked GWAS. From a network with the WTCCC2 MS associations omitted, we predicted probabilities of association for all potentially novel genes. The 37 novel genes identified by the WTCCC2 GWAS were considered positives, and the resulting performance was plotted. The ROC (A) and precision-recall (B) curves show performance, with AUCs in line with the testing performance across all diseases (Figure 2). A prediction threshold (black cross) that resulted in high performance was selected as the discovery threshold for further analysis. As the classification threshold decreases along the precision-recall curve, the advent of each true positive is denoted by its gene symbol.

### Prioritizing statistical candidates with network-based predictions identifies novel multiple sclerosis genes

Finally, we designed a framework for discovering and validating novel MS genes that incorporates our network-based predictions. Meta2.5 is a meta-analysis of all MS GWAS prior to the WTCCC2 study [13]. We calculated genewise p-values for Meta2.5 using VEGAS [14] and observed a large enrichment in nominally significant (*p* < 0.05) genes, suggesting multiple potential associations (Figure S8). We combined this set of experimental candidates with the top predictions from the pre-WTCCC2 network to discover genes with both strong statistical and biological evidence of association. To ensure novelty, we excluded genes from GWAS-established MS loci and the extended MHC region. We chose a threshold (Table S3) for network-based predictions that performed well in prioritizing the genes identified by WTCCC2 (Figure 5).

This strategy discovered four genes, three of which—*JAK2*, *REL*, *RUNX3* —achieved Bonferroni validation on VEGAS-converted WTCCC2 p-values (Table 4). The probability of the observed validation rate occurring under random prioritization is 0.01 (Table S3), demonstrating that incorporating our network-based predictions as a prior increased study power. *JAK2* displays overexpression in MS-affected Th17 cells [15] and was implicated in an interactome-based prioritization of GWAS [2]. *RUNX3*, a transcription factor influencing T lymphocyte development, has been associated with celiac disease [16] and ankylosing spondylitis [17] and was hypermethylated in systemic lupus erythematosus patients [18]. The region containing *REL* was uncovered in a recent MS ImmunoChip-based study with 14,498 cases [19, p. S40]. For the gene-dense region containing *REL*, the ImmunoChip study reported a long non-coding RNA, *LINC01185*, overlapping the lead-SNP, rs842639. However, since greater than 80% of the genome shows evidence of transcription [20], the probability of incidental overlap with long non-coding RNA is high. *REL*, however, is an essential transcription factor for lymphocyte development [21] and plays a critical role in autoimmune inflammation [22]. Hence, gene prioritization through integrative analyses offers not only to streamline loci discovery but also subsequent causal gene identification.

**Table 4.** Multiple sclerosis gene discovery. Four genes showed nominal statistical evidence of association (Meta2.5 column) and exceeded the network prediction threshold (HNLP column). Three genes achieved Bonferroni validation (bold) in an independent GWAS (WTCCC2 column).

| Gene | Meta2.5 | HNLP | WTCCC2 |
| --- | --- | --- | --- |
| JAK2 | 0.047 | 0.102 | **0.0015** |
| REL | 0.001 | 0.040 | **0.0003** |
| SH2B3 | 0.012 | 0.034 | 0.0130 |
| RUNX3 | 0.016 | 0.025 | **0.0073** |

## Discussion

In this work, we developed a framework to predict the probability that each protein-coding gene is associated with each of 29 complex diseases. Our predictions draw on a diverse set of pathogenically-relevant relationships encoded in a heterogeneous network. The predictions successfully prioritized associations hidden from the network. Using MS as a representative example, we were able to combine our predictions with statistical evidence of association to increase study power and identify three novel susceptibility genes in this disease. The disease-specific performance (measured by the AUROC) for MS was exceeded by twelve other diseases suggesting that our predictions have broad applicability for prioritizing genetic association analyses. Prioritization can range from a genome-wide scale to a single loci where this approach can highlight the causal gene from several candidates within the same association block. For researchers focused on a specific disease, these predictions can be used to propose genes for experimental investigation. Inversely, researchers focused on a specific gene can use this resource to find suggestions for relevant complex disease phenotypes.

Most previous explorations of the factors underlying pathogenicity have focused on a single domain such as tissue-specificity [23], protein interactions [24], pathways [1], or disease similarity [25]. The method presented here integrates disparate data sources, learns their importance, and unifies them under a common framework enabling comparison. Therefore, we can conclude that perturbation gene sets—the core of our top-performing feature—are an underutilized resource for disease-associated gene prioritization. Not only did perturbations encompass other set-based gene categorizations, but they greatly outperformed features based on protein interactions, pathways, and tissue-specificity, which form the basis of several prominent prioritization techniques. In addition to characterizing the overall importance of each feature, our online prediction browser visually decomposes an individual prediction into its components.

We observed a prominent influence of pleiotropy, consistent with previous studies that identified pervasive overlap of susceptibility loci across complex diseases [26], especially those of autoimmune nature [27]. Since many existing prioritization techniques are agnostic to the compendia of GWAS associations, they fail to adequately leverage pleiotropy. Unlike approaches initiated from a user-provided gene list, our study only provides predictions for 29 diseases. By not relying on user-provided input, our predictions can serve as independent priors for future analyses. By predicting probabilities, we provide an extensible and interpretable assessment of association that circumvents the limitations inherent to frequentist analyses [28]. Many approaches return no assessment for the majority of genes which fall outside of their set of predicted positives. Here, we overcome this issue and provide a comprehensive and genome-wide output by returning a probability of association for each protein-coding gene.

High-throughput biological data is frequently noisy and incomplete [29]. Combining orthogonal resources can help overcome these issues. Accordingly, we found that our integrative model outperformed any individual domain. While this method has shown encouraging performance, some limitations are worth noticing. For example, many biological networks preferentially cover well-studied vicinities [30]. Knowledge biases that span multiple presumably-orthogonal resources could diminish the benefits of integration. Here, several of the literature-derived domains were removed by the lasso suggesting redundancy. Biases in network completeness can also lead to high-quality predictions for well-studied vicinities and low-quality predictions for poorly-studied vicinities. The permutation analysis provided evidence of this disparity: edge-specificity was critical for top predictions yet only moderately beneficial for the remainder of predictions. Subsequently, we caution users to avoid overinterpreting predictions for poorly-characterized genes. To help place predictions in context, the online browser provides a gene’s mean prediction across all diseases and a disease’s mean prediction across all genes. As more systematic and unbiased resources become available [29], high-quality predictions will be possible for a higher percentage of network vicinities.

We reason that the desirable qualities of our predictions are the consequence of the heterogenous network edge prediction methodology. The approach is versatile (most biological phenomena are decomposable into entities connected by relationships), scalable (no theoretical limit to metagraph complexity or graph size), and efficient (low marginal cost to including an additional network component). We have extended the previous metapath-based framework set forth by *PathPredict* [7], by: 1) incorporating regularization allowing coefficient estimation for more features without overfitting; 2) designing a framework for predicting a metaedge that is included in the network; 3) developing an improved metric for assessing path specificity; and 4) implementing a degree-preserving permutation. Metapath-based heterogeneous network edge prediction provides a powerful new platform for bioinformatic discovery.

## Methods

### Heterogeneous networks

We created a general framework and open source software package for representing heterogeneous networks. Like traditional graphs, heterogeneous networks consist of nodes connected by edges, except that an additional meta layer defines type. Node type signifies the kind of entity encoded, whereas edge type signifies the kind of relationship encoded. Edge types are comprised of a source node type, target node type, kind (to differentiate between multiple edge types connecting the same node types), and direction (allowing for both directed and undirected edge types). The user defines these types and annotates each node and edge, upon creation, with its corresponding type. The meta layer itself can be represented as a graph consisting of node types connected by edge types. When referring to this graph of types, we use the prefix ‘meta’. Metagraphs—called schemas in previous work [6, 7]—consist of metanodes connected by metaedges. In a heterogeneous network, each path, a series of edges with common intermediary nodes, corresponds to a metapath representing the type of path. A path’s metapath is the series of metaedges corresponding to that path’s edges. The possible metapaths within a heterogeneous network can be enumerated by traversing the metagraph. We implemented this framework as an object-oriented data structure in python and named the resulting package *hetio*. Users are free to browse, use, or contribute to the software, through the online repository (http://github.com/dhimmel/hetio).

## Network construction

Protein-coding genes were extracted from the HGNC database [31]. Resources were mapped to HGNC terms via gene symbol (ambiguous symbols were resolved in the order: approved, previous, synonyms) or Entrez identifiers. Disease nodes were taken from the Disease Ontology (DO) [32]. Due to the limited number of diseases with GWAS, relevant disease references were manually mapped to the DO. Tissues were taken from the BRENDA Tissue Ontology (BTO) [33]. Only tissues with profiled expression were included enabling manual mapping. Nodes for the 14 MSigDB metanodes were directly imported from the Molecular Signature Database version 4.0 [34]. MSigDB collections that were supersets of other collections were excluded. Diseases were classified manually into 10 categories according to pathophysiology. The ‘idiopathic’ and ‘unspecific’ categories were not included as pathophysiology nodes, since they do not signify meaningful similarities between member diseases.

### Association processing

Disease-gene associations were extracted from the GWAS Catalog [8], a compilation of GWAS associations where *p* < *10*^−5^. First, associations were segregated by disease. GWAS Catalog phenotypes were converted to Experimental Factor Ontology (EFO) terms using mappings produced by the European Bioinformatics Institute. Associations mapping to multiple EFO terms were excluded to eliminate cross-phenotype studies. We manually mapped EFO to DO terms (now included in the DO as cross-references) and annotated each DO term with its associations.

Associations were classified as either high or low-confidence, where exceeding two thresholds granted high-confidence status. First, *p* ≤ 5 × 10^−8^ corresponding to *p* ≤ 0.05 after Bonferroni adjustment for one million comparisons (an approximate upper bound for the number of independent SNPs evaluated by most GWAS). Second, a minimum sample size (counting both cases and controls) of 1,000 was required, since studies below this size are underpowered [35]—i.e. any discovered associations are more likely than not to be false—for the majority of true effect size distributions commonly assumed to underlie complex disease etiology [28].

Lead-SNP were assigned windows—regions wherein the causal SNPs are assumed to lie—retrieved from the DAPPLE server [4]. Windows were calculated for each lead-SNP by finding the furthest upstream and downstream SNPs where *r*^2^ > 0.5 and extending outwards to the next recombination hotspot. Associations were ordered by confidence, sorting on following criteria: high/low confidence, p-value (low to high), and recency. In order of confidence, associations were overlapped by their windows into disease-specific loci. By organizing associations into loci, associations from multiple studies tagging the same underlying signal were condensed. A locus was classified as high-confidence if any of its composite associations were high-confidence and low-confidence otherwise.

For each disease-specific loci, we attempted to identify a primary gene. The primary gene was resolved in the following order: 1) the mode author-reported gene; 2) the containing gene for an intragenic lead-SNP; 3) the mode author-reported gene for an intragenic lead-SNP (in the case of overlapping genes); 4) the mode author-reported gene of the most proximal up and downstream genes. Steps 2–4 were repeated on each association composing the loci, in order of confidence, until a single gene resolved as primary. Loci where ambiguity was unresolvable or where no genes were returned did not receive a primary gene. All non-primary genes—genes that were author-reported, overlapping the lead-SNP, or immediately up or downstream from the lead-SNP—were considered secondary.

Accordingly, four categories of processed associations were created: high-confidence primary, high-confidence secondary, low-confidence primary, and low-confidence secondary. We assume that our primary gene annotation for each loci represents the single causal gene responsible for the association. To investigate the validity of this assumption, we evaluated the performance of our predictions separately using each category of association as positives (Figure S7). For both confidence levels, primary associations outperformed secondary associations suggesting our method succeeded at categorizing causal genes as primary. However, for high-confidence secondary associations, the AUROC equaled 0.74, which could result from multiple causal genes per loci or categorizing sole causal genes as secondary. The performance decline from high to low confidence associations was severe, pointing to a preponderance of falsely identified loci in the GWAS Catalog when *p* > 5 × 10^−8^ or sample size drops below 1000.

### Protein interactions

Physical protein-protein interactions were extracted from iRefIndex 12.0, a compilation of 15 primary interaction databases [36]. The iRefIndex was processed with ppiTrim to convert proteins to genes, remove protein complexes, and condense duplicated entries [37].

### Tissue-specific gene expression

Tissue-specific gene expression levels were extracted from the GNF Gene Expression Atlas [38]. Starting with the GCRMA-normalized and multisample-averaged expression values, 44,775 probes were converted to 16,466 HGNC genes and 84 tissues were manually mapped and converted to 77 BTO terms. For both conversions, the geometric mean was used to average expression values. The log base 10 of expression value was used as the threshold criteria for *GeT* edge inclusion.

### Disease localization

Disease localization was calculated for the 77 tissues with expression profiles. Literature co-occurrence was used to assess whether a tissue is affected by a disease. We used CoPub 5.0 to extract R-scaled scores between tissues and diseases measuring whether two terms occurred together in Medline abstracts more than would be expected by chance [39]. DO terms for diseases with GWAS and BTO tissues with expression profiles were manually mapped to the ‘biological identifier’ terminology used by CoPub. The R-scaled score was used as the threshold criteria for *TlD* edge inclusion.

## Feature computation metrics

The simplest metapath-based metric is path count (*PC*): the number of paths, of a specified metapath, between a source and target node. However, *PC* does not adjust for the extent of graph connectivity along the path. Paths traversing high-degree nodes will account for a large portion of the *PC*, despite high-degree nodes frequently representing a biologically broad or vague entity with little informativeness. The previous work evaluated several metrics that include a *PC* denominator to adjust for connectivity and reported that normalized path count (*NPC*) performed best [7]. The denominator for *NPC* equals the number of paths from the source to any target plus the number of paths from any target to the source. We adopt the any source/target concept to compute the two *GaD* features. However, dividing the *PC* by a denominator is flawed because each path composing the PC deserves a distinct degree adjustment. If two paths—one traversing only high-degree nodes and one traversing only low-degree nodes—compose the *PC*, the network surrounding the high-degree path will monopolize the *NPC* denominator and overwhelm the contribution of the low-degree path despite its specificity. Therefore, we developed the degree-weighted path count (*DWPC*) which individually downweights each path between a source and target node. Each path receives a path-degree product (*PDP*) calculated by: 1) extracting all metaedge-specific degrees along the path (each edge composing the path contributes two degrees); 2) raising each degree to the *−w* power, where *w ≥* 0 and is called the damping exponent; 3) multiplying all exponentiated degrees to yield the *PDP*. The *DWPC* equals the sum of *PDPs*. See Figure S2C–D for a visual and algebraic description of the *DWPC*.

## Machine learning approach

Regularized logistic regression requires a parameter, *λ*, setting the strength of regularization. We optimized *λ* separately for each model fit. Using 10-fold cross-validation and the “one-standard-error” rule to choose the optimal *λ* from deviance, we adopted a conservative approach designed to prevent overfitting [40].

On the training set of gene-disease pairs, we optimized the elastic net mixing parameter (*α*), the *DWPC* damping exponent (*w*), and two edge inclusion thresholds. First, we optimized *α* and *w* on the 20 features whose metapaths did not include threshold-dependent metaedges. For each combination of *α* and *w*, we calculated average testing AUROC using 20-fold cross-validation repeated for 10 randomized partitionings. After setting *α* and *w* (Figure S3A), we jointly optimized the two edge-inclusion thresholds using the AUROC for the *GeTlD* feature, whose metapath is composed from the two edges requiring thresholds (Figure S3B).

## Degree-preserving permutation

Starting from the complete network, a permuted network was created by swapping edges separately for each metaedge. Edge swaps were performed by switching the target nodes for two randomly selecting edges [41]. For each metaedge, the number of attempted swaps was ten times the corresponding edge count. We adopted a Markov Chain strategy where additional rounds of permutation were initiated from the most-recently permuted network [41]. A training network was generated from the first permuted network by masking 25% of the associations for testing. Testing performance for the permuted training network model is shown by Figure S4. When contrasting this performance with the unpermuted-network model, we employed the Condensed-ROC curve to magnify the importance of top predictions [42]. Using the exponential transformation with a magnification factor of 460—the value which maps a FPR of 0.01 to 0.99—we concentrated on the top 1% of predictions (Figure S4C). A one-sided unpaired DeLong test [43] was used to assess whether feature-specific AUROCs from the complete network exceeded those from the first permuted network (Table S2).

## Multiple sclerosis gene discovery

We excluded 588 genes from the discovery phase of the multiple sclerosis analysis. First we excluded genes in the extended MHC region (spanning from *SCGN* to *SYNGAP1* on chromosome 6 [44]) due to the complex pattern of linkage characterizing this region containing several highly-penetrant MS-risk alleles [12]. Second, we excluded putative MS genes: high-confidence primary genes from the GWAS Catalog and reported genes for the WTCCC2-replicated loci. We omitted genes in linkage disequilibrium with the putative genes by excluding: 1) consecutive sequences of nominally significant genes (using the WTCCC2-VEGAS p-values) that included a putative gene; and 2) high-confidence secondary genes from the GWAS catalog. Post exclusion, 1211 genes were nominally significant in Meta2.5, four of which exceeded the network-based discovery threshold. Using a hypergeometric test for overrepresentation, we calculated the probability of randomly selecting 4 of the 1211 genes and Bonferroni validating at least 3 of the 4 on WTCCC2 (Table S3).

## Data availability

See Datasets S1–10 for the supporting data. The website provides additional resources (http://het.io/disease-genes/downloads/) as well as an interface for browsing results (http://het.io/disease-genes/browse/). Project related code is available from the github repository (http://github.com/dhimmel/hetio).

## Ethics Statement

This study was approved by the UCSF institutional review board on human subjects under protocol #10-00104.

## Supporting Figure Legends

**Figure S1.**
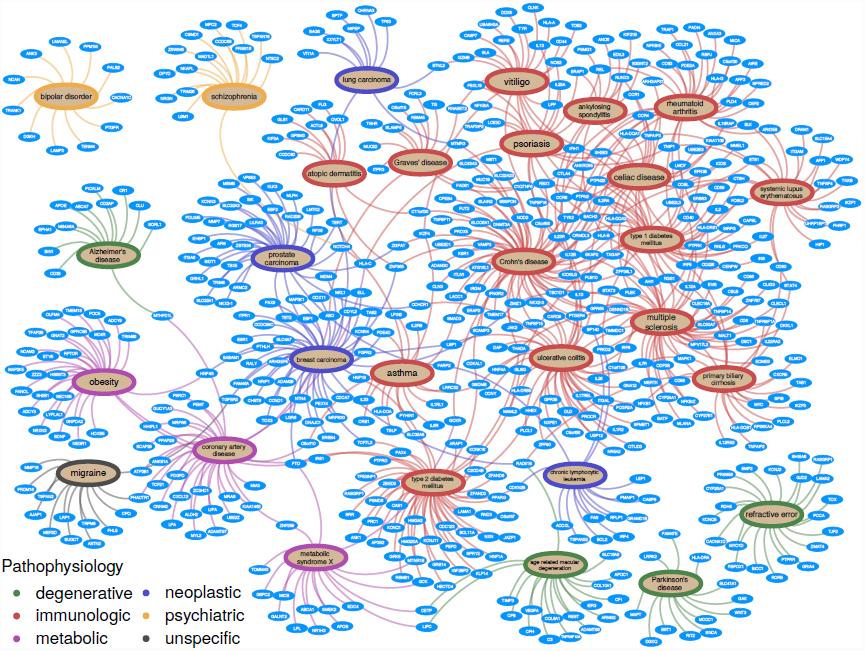
Bipartite network of gene-disease associations. Gene-disease associations were extracted from the GWAS Catalog. Here we show the 698 high-confidence primary associations for the 29 diseases with at least 10 associations. Diseases (large nodes) and their incident edges are colored according to disease pathophysiology. The network highlights pervasive pleiotropy as well as the overlap of susceptibility genes among autoimmune diseases.

**Figure S2.**
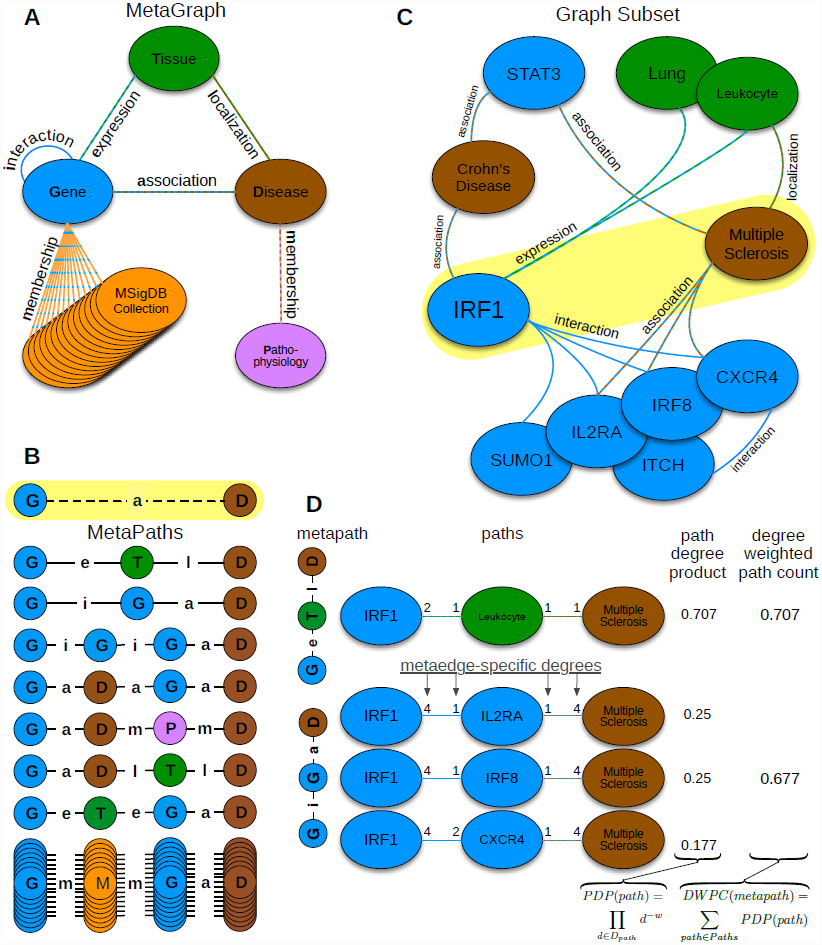
Heterogeneous network edge prediction methodology. A) We constructed the network according to a schema, called a metagraph, which is composed of metanodes (node types) and metaedges (edge types). B) The network topology connecting a gene and disease node is measured along metapaths (types of paths). Starting on Gene and ending on Disease, all metapaths length three or less are computed by traversing the metagraph. C) A hypothetical graph subset showing select nodes and edges surrounding *IRF1* and multiple sclerosis. To characterize this relationship, features are computed that measure the prevalence of a specific metapath between IRF1 and multiple sclerosis. D) Two features (for the *GeTlD* and *GiGaD* metapaths) are calculated to describe the relationship between IRF1 and multiple sclerosis. The metric underlying the features is degree-weighted path count (*DWPC*). First, for the specified metapath, all paths are extracted from the network. Next, each path receives a path-degree product measuring its specificity (calculated from node-degrees along the path, *D*_*path*_). This step requires a damping exponent (here *w* = 0.5), which adjusts how severely high-degree paths are downweighted. Finally, the path-degree products are summed to produce the *DWPC*.

**Figure S3.**
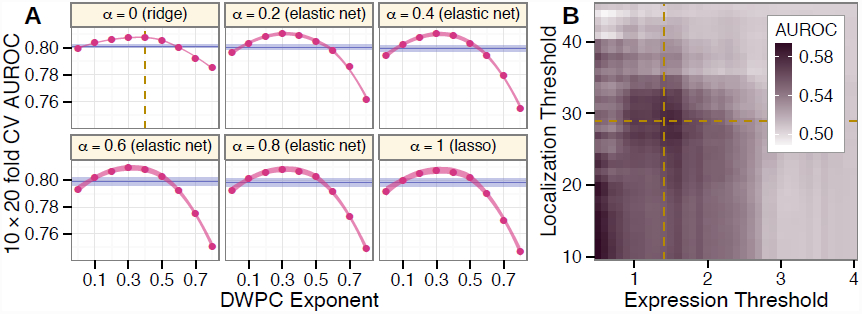
Parameter optimization. Using the training network, optimal parameter values (yellow dashed lines) were chosen. A) Using average cross-validated AUROC to assess performance, six elastic net mixing parameters were evaluated. For each mixing parameter value *α*, 10 feature metrics were evaluated: the *DWPC* for 9 weighting exponents (*w*, magenta with a 99.99% loess confidence band) and the *NPC* (violet with a 99.99% confidence interval). The *DWPC* with *w* = 0.4 outperformed the *NPC*, the best metric from previous work, as well as the path count which equals the *DWPC* when *w* = 0. Performance variability was minimized when *α* = 0. B) Edge-inclusion thresholds for two metaedges were jointly optimized. Expression threshold refers to the minimum microarray intensity required for a tissue-specific expression (*GeT*) edge. Localization threshold refers to the minimum literature co-occurrence score required for a disease localization (*TlD*) edge. Treating the *DWPC* (*w* = 0.4) for the *GeTlD* metapath as a classifier, the AUROC was calculated at each pairwise threshold combination. The optimal thresholds were chosen as the center of a stable, high-performing, and computationally-feasible section of the solution space.

**Figure S4.**
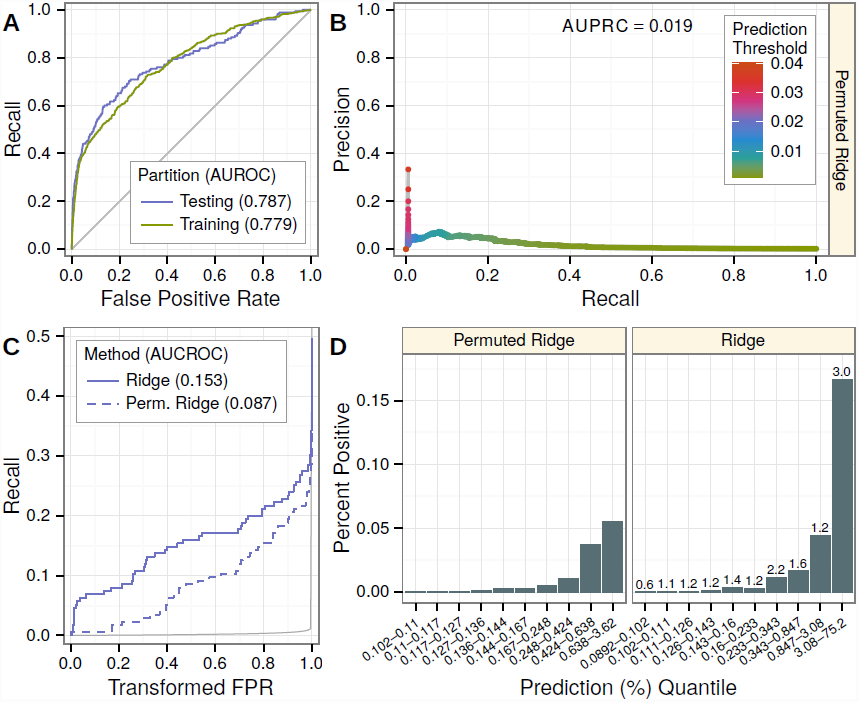
Performance of the degree-preserving permutation. Testing performance is contrasted between ridge models for the permuted-network and unpermuted-network. A) Testing and training ROC curves for the permuted-network model. B) Testing precision-recall curve for the permuted-network model. C) Testing CROC curves for the permuted-network and unpermuted-network models. The FPR has been scaled to focus on the first 1% placing greater emphasis on top predictions. While both models vastly outperform random (grey line), the unpermuted-network model provides far superior top predictions. D) For both networks, gene-disease pairs were stratified by deciles of the predicted probabilities for positives. For each strata, the percent of positive pairs (precision) is plotted. The fold change over permuted is denoted for the unpermuted deciles.

**Figure S5.**
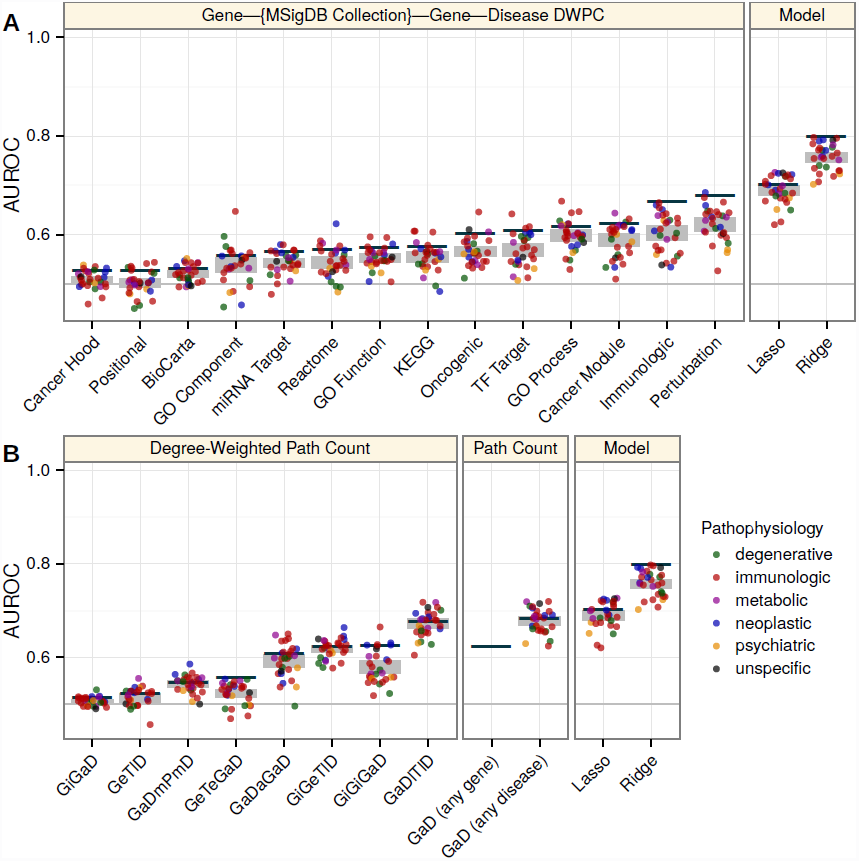
Disease, feature, and model-specific performance across permuted-network models. Disease, feature, and model-specific AUROCs were calculated separately for each of 5 permuted networks and averaged. The figure is analogous to Figure 4, except all measures refer to permuted-network performance. Disease-specific performance tends towards the mean, as disease-specific information has been altered by permutation. For features ending with an association (*GaD*) metaedge, global performance exceeds disease-specific performance. These features capture disease polygenicity, which improves the ranking of gene-disease pairs only if multiple diseases are included. Performance of the lasso model is affected, since the signals become too weak and few features survive regularization.

**Figure S6.**
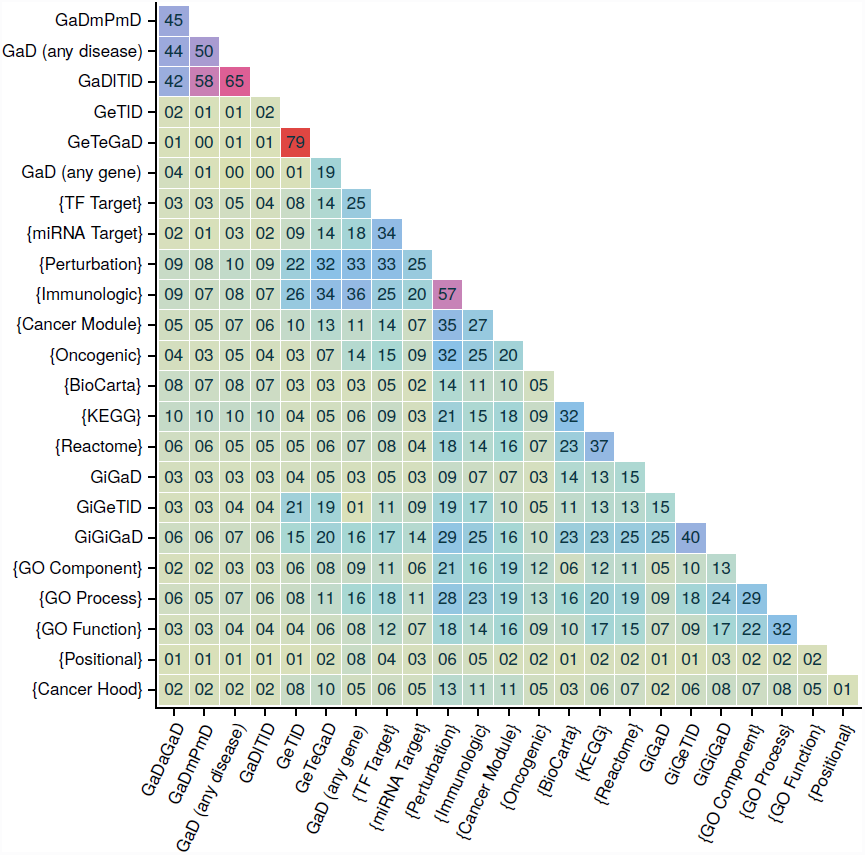
Pairwise feature correlation. Pearson’s correlation coefficients (shown by color and as a percent) were calculated for all pairwise feature combinations. Features were ordered using Ward’s hierarchical clustering. Moderate collinearity is pervasive across features. The four pleiotropy-focused features form a tight cluster (top left). *Perturbations* and *Immunologic signatures* are correlated with many other features, including several other MSigDB features.

**Figure S7.**
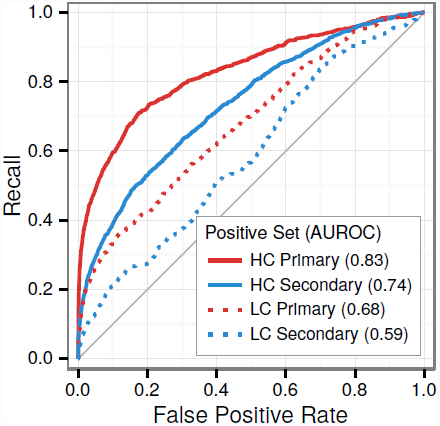
Performance of the predictions on the four categories of associations. Keeping unassociated gene-disease pairs as negatives, ROC curves were calculated separately for each category of association as positives. Predictions from the complete-network ridge model were used as the classifier. For both high and low-confidence associations, primary gene annotations received higher predictions than secondary gene annotations. High-confidence associations received considerably higher predictions than low-confidence associations suggesting a high frequency of false positives amongst low-confidence associations.

**Figure S8.**
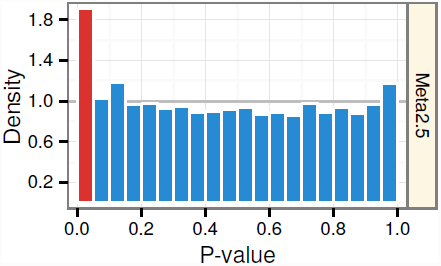
Excess of nominally significant genewise p-values in Meta2.5. The histogram of genewise p-values from Meta2.5, a meta-analysis of multiple sclerosis GWAS preceding the WTCCC2 study. If no associations are present, uniformly distributed p-values (grey line) would be expected. Instead, we observed an excess of nominally significant genes (*p* ≤ 0.05, red) indicating a set of genes likely enriched for true associations.

**Table S1.** Features. The 24 features computed for each gene-disease pair and the aspect of network topology described.

| **Path Count** | **Measures the number of ...** |
| --- | --- |
| GaD (any disease) | diseases that the source gene is associated with, ignoring the association with the target disease if present. |
| GaD (any gene) | genes that the target disease is associated with, ignoring the association with the source gene if present. |
| **DWPC** | **Measures the extent that ...** |
| GeTID | the source gene is expressed in tissues affected by target disease. |
| GiGaD | genes associated with the target disease interact with the source gene. |
| GiGiGaD | genes associated with the target disease interact with genes that interact with the source gene. |
| GaDaGaD | genes associated with the same diseases as the source gene are associated with the target disease. |
| GaDmPmD | diseases with the same pathophysiology as the target disease are associated with the source gene. |
| GaDITID | diseases affecting the same tissues as the target disease are associated with the source gene. |
| GeTeGaD | genes expressed in the same tissues as the source gene are associated with the target disease. |
| GiGeTID | genes interacting with the source gene are expressed in tissues that are affected by the target disease. |
| {Positional} | genes located in the same cytogenetic band as the source gene are associated with the target disease. |
| {Perturbation} | genes belonging to the same perturbation signatures as the source gene are associated with the target disease. |
| {BioCarta} | genes involved in the same BioCarta pathways as the source gene are associated with the target disease. |
| {KEGG} | genes involved in the same KEGG pathways as the source gene are associated with the target disease. |
| {Reactome} | genes involved in the same Reactome pathways as the source gene are associated with the target disease. |
| {miRNA Target} | genes sharing 3'-UTR microRNA binding motifs with the source gene are associated with the target disease. |
| {TF Target} | genes sharing transcription factor binding sites with the source gene are associated with the target disease. |
| {Cancer Hood} | genes present in the same expression neighborhoods of cancer-related genes as the source gene are associated with the target disease. |
| {Cancer Module} | genes belonging to the same cancer modules as the source gene are associated with the target disease. |
| {GO Process} | genes participating in the same GO Biological Processes as the source gene are associated with the target disease. |
| {GO Component} | genes belonging to the same GO Cellular Components as the source gene are associated with the target disease. |
| {GO Function} | genes contributing to the same GO Molecular Functions as the source gene are associated with the target disease. |
| {Oncogenic} | genes belonging to the same cancer-dysregulated cellular pathways as the source gene are associated with the target disease. |
| {Immunologic} | genes belonging to the same immunologic signatures as the source gene are associated with the target disease. |

**Table S2.** Feature-specific performance before and after network permutation. Ten features (bold) showed a significant (*p* < 0.05, one-sided DeLong test) decrease in performance.

| Feature | AUROC | p-AUROC | p-value |
| --- | --- | --- | --- |
| **GaDmPmD** | 0.643 | 0.547 | $1.6 \times 10^{-20}$ |
| **GiGaD** | 0.558 | 0.514 | $2.1 \times 10^{-9}$ |
| **{Perturbation}** | 0.740 | 0.667 | $2.3 \times 10^{-7}$ |
| **GeTID** | 0.573 | 0.518 | $1.8 \times 10^{-5}$ |
| **{KEGG}** | 0.613 | 0.566 | $5.6 \times 10^{-5}$ |
| **GaDaGaD** | 0.633 | 0.592 | $1.5 \times 10^{-4}$ |
| **{BioCarta}** | 0.548 | 0.526 | 0.001 |
| **{Reactome}** | 0.599 | 0.562 | 0.002 |
| **{Immunologic}** | 0.703 | 0.665 | 0.006 |
| **GiGiGaD** | 0.646 | 0.621 | 0.05 |
| {Cancer Module} | 0.629 | 0.612 | 0.11 |
| GeTeGaD | 0.570 | 0.554 | 0.14 |
| {TF Target} | 0.612 | 0.596 | 0.16 |
| {GO Component} | 0.560 | 0.547 | 0.17 |
| {Positional} | 0.529 | 0.520 | 0.19 |
| GiGeTID | 0.628 | 0.616 | 0.20 |
| {GO Process} | 0.626 | 0.617 | 0.26 |
| GaD (any disease) | 0.683 | 0.676 | 0.30 |
| {GO Function} | 0.577 | 0.571 | 0.31 |
| {Cancer Hood} | 0.533 | 0.532 | 0.45 |
| GaD (any gene) | 0.620 | 0.620 | 0.49 |
| GaDITID | 0.674 | 0.674 | 0.49 |
| {Oncogenic} | 0.601 | 0.604 | 0.60 |
| {miRNA Target} | 0.562 | 0.569 | 0.72 |

**Table S3.** Multiple sclerosis gene discovery statistics. The upper section details the high-performing network prediction threshold. The lower section details the hypergeometric test for overrepresentation of validating genes.

|  | Value |
| --- | --- |
| **Prediction Threshold** | 0.024 |
| **False Positive Rate** | 0.001 |
| **Recall** | 0.108 |
| **Precision** | 0.133 |
| **Lift** | 68.4 |
| **Novel & Meta2.5-nominal Total** | 1211 |
| **Discovered** | 4 |
| **Bonferroni Cutoff** | 0.0125 |
| **Discovered < Bonferroni** | 3 |
| **Total < Bonferroni** | 199 |
| **Replication $p$-value** | 0.015 |

## Supporting Data

**Dataset S1. Predictions.** Predicted probabilities of association between all genes (rows) and diseases (columns).

**Dataset S2. Features.** The features (columns) computed for each gene-disease pair (rows). Column names with beginning with ‘XB’ refer to standardized features.

**Dataset S3. Serialized network.** A JSON formatted text file storing the complete network. The top level is an JSON object with four pairs (metanodes, metaedges, nodes, edges). The value for each pair is a JSON array containing the corresponding items.

**Dataset S4. Processed GWAS Catalog Loci.** Loci-disease associations. The file includes the gene resolution information for each loci including the studies and SNPs underlying the association.

**Dataset S5. Gene-Disease Associations.** All gene-disease associations extracted from the GWAS catalog for the four categories of association.

**Dataset S6. Disease Ontology Modifications.** Ten DO terms that appeared in the GWAS Catalog were redundant with other terms. Seven were removed. Three were merged with recipient terms by removing the term and transferring the associations.

**Dataset S7. Tissue-specific gene expression.** A processed version of the GNF BodyMap providing a gene’s (row, HGNC symbols) expression value for each of 77 tissues (columns, BRENDA Tissue Ontology IDs).

**Dataset S8. Disease localization.** Literature co-occurrence scores between diseases and tissues computed using CoPub 5.0.

**Dataset S9. Terminology Mappings.** All mappings that were manually performed. Specifically, tissue and disease mappings to CoPub ’Biologic Identifiers’, tissue mappings to GNF BodyMap samples, disease mappings to the EFO terms appearing in the GWAS Catalog, and disease pathophysiologies.

**Dataset S10. Multiple Sclerosis Analysis.** For each gene (row), the genewise Meta2.5 and WTCCC2 p-values and network-based predictions are reported.

**Dataset S11. Vector Images.** PDF formatted versions of the figures.

